# A mechanism of uncompetitive inhibition of the serotonin transporter

**DOI:** 10.1101/2022.08.11.503588

**Authors:** Shreyas Bhat, Ali El-Kasaby, Ameya Kasture, Danila Boytsov, Julian B. Reichelt, Thomas Hummel, Sonja Sucic, Christian Pifl, Michael Freissmuth, Walter Sandtner

**Affiliations:** Institute of Pharmacology and the Gaston H. Glock Research Laboratories for Exploratory Drug Development, Center of Physiology and Pharmacology, Medical University of Vienna, Vienna, Austria; Department of Neurobiology, University of Vienna, Vienna, Austria; Center for Brain Research, Medical University of Vienna, Vienna, Austria

**Keywords:** serotonin transporter, N-formyl-1,3-bis (3,4-methylenedioxyphenyl)-prop-2-yl-amine – ECSI#6, misfolding, pharmacochaperoning, cocaine, noribogaine

## Abstract

The serotonin transporter (SERT/SLC6A4) is arguably the most extensively studied solute carrier (SLC). During its eponymous action - i.e., the retrieval of serotonin from the extracellular space - SERT undergoes a conformational cycle. Typical inhibitors (antidepressant drugs and cocaine), partial and full substrates (amphetamines and their derivatives) and atypical inhibitors (ibogaine analogues) bind preferentially to different states in this cycle. This results in competitive or non-competitive transport inhibition. Here, we explored the action of N-formyl-1,3-bis (3,4-methylenedioxyphenyl)-prop-2-yl-amine (ECSI#6) on SERT: inhibition of serotonin uptake by ECSI#6 was enhanced with increasing serotonin concentration. Conversely, the K_M_ for serotonin was lowered by augmenting ECSI#6. ECSI#6 bound with low affinity to the outward-facing state of SERT but with increased affinity to a potassium-bound state. Electrophysiological recordings showed that ECSI#6 preferentially interacted with the inward-facing state. Kinetic modeling recapitulated the experimental data and verified that uncompetitive inhibition arose from preferential binding of ECSI#6 to the K^+^-bound, inward-facing conformation of SERT. This binding mode predicted a pharmacochaperoning action of ECSI#6, which was confirmed by examining its effect on the folding-deficient mutant SERT-PG^601,602^AA: pre-incubation of HEK293 cells with ECSI#6 restored export of SERT-PG^601,602^AA from the endoplasmic reticulum and substrate transport. Similarly, in transgenic flies, administration of ECSI#6 promoted delivery of SERT-PG^601,602^AA to the presynaptic specialization of serotonergic neurons. To the best of our knowledge, ECSI#6 is the first example of an uncompetitive SLC inhibitor. Pharmacochaperones endowed with the binding mode of ECSI#6 are attractive, because they can rescue misfolded transporters at concentrations, which cause modest transport inhibition.

## Introduction

The transporters for the monoamines serotonin (SERT), dopamine (DAT), and norepinephrine (NET) are members of the solute carrier-6 (SLC6) family [1]. The uptake of their eponymous substrates is fueled by the electrochemical gradient of sodium; in addition, DAT and NET can harvest the membrane potential, while SERT utilizes the potassium gradient [2]. These driving forces support a remarkable concentrative power, which allows for effective removal of the cognate substrates from the extracellular space. SERT, NET and DAT are closely related. Accordingly, their pharmacology shows a continuum from compounds, which bind to all three transporters (e.g. cocaine), to drugs with exquisite selectivity (e.g. selective serotonin reuptake inhibitors, SSRIs). Monoamine transporters are targets for both, therapeutically relevant drugs (e.g. antidepressants) and for illicit substances. Accordingly, the chemical space of ligands has been extensively studied resulting in a very rich pharmacology [3]. Originally, ligands for monoamine transporters have been classified as either nontransportable inhibitors or exogenous substrates/releasers. Inhibitors bind to and block reuptake through the transporter (e.g., cocaine, tricyclic antidepressants, SSRIs). In contrast, exogenous substrates/releasers (e.g., amphetamines) induce efflux of the endogenous monoamine by switching the transporter from the physiological forward transport mode into an exchange mode [3,4].

More recently, the prevailing dichotomous classification was challenged by the discovery of compounds, which act as partial substrates and/or atypical inhibitors of monoamine transporters [5–10]. Partial substrates do not elicit reverse transport to the same extent as full releasers [5,7]. In fact, there is a continuum, which ranges from full releasers over partial releasers to atypical inhibitors: minor modifications of the amphetamine- or cathinone-based structures suffice to reduce the efficacy of release and to eventually produce atypical inhibitors or allosteric modulators [7,9]. Typical inhibitors bind to the outward facing conformation of the monoamine transporters. In contrast, atypical inhibitors trap the transporter in distinct conformational states, which are visited during the transport cycle [11]. Numerous disease-associated mutations have been identified in neurotransmitter transporters, which result in misfolding of the protein [12]. Atypical inhibitors and partial substrates are of interest, because they can act as pharmacochaperones and correct the folding defect [11,13]. Here, we studied the inhibition of SERT by the amphetamine derivative ECSI#6 [N-formyl-1,3-bis (3,4-methylenedioxyphenyl)-prop-2-yl-amine], which was identified as a synthesis byproduct of 3,4-methylenedioxy-methamphetamine (MDMA,”ecstasy”) [14]. Our experiments show that ECSI#6 is, to the best of our knowledge, the first reported uncompetitive inhibitor of SERT. This mode of inhibition was accounted for by preferential binding of ECSI#6 to the inward-facing state. Finally, consistent with this binding mode, ECSI#6 also acted as a pharmacochaperone: cellular preincubation with ECSI#6 restored export from the endoplasmic reticulum and substrate transport by a misfolded SERT variant.

## Material and Methods

### Cell Culture and Materials

HEK293 cells were cultured in Dulbecco’s modified Eagle’s medium (DMEM) supplemented with 10% heat-inactivated fetal bovine serum (FBS), 0.6 mg. L^-1^ penicillin and 1 mg. L^-1^ streptomycin and 5 mg. L^-1^ plasmocin. These cells were transfected by combining either YFP-tagged WT-SERT or YFP-tagged SERT-PG^601,602^AA with PEI (linear 25 kDa polyethylenimine; Santa Cruz, SC-360988A) at a ratio of 1:3 (w/w) in serum-free DMEM. HEK293 cells stably expressing YFP-tagged SERT, which were used for uptake experiments, were cultured in medium supplemented with 100□mg.L ^1^ geneticin (G418) for clonal selection. HEK293 cells expressing GFP-tagged SERT, which were used for binding experiments and electrophysiology, were cultured in medium supplemented with 150□mg. L^-1^ of zeocin and 6□mg. L^-1^ blasticidin. The expression of GFP-SERT was induced by 1 mg. L^-1^ tetracycline 24 h prior to either membrane preparation or electrophysiology. All antibiotics were purchased from InvivoGen (USA). For the pharmacochaperoning experiments, noribogaine was purchased from Cfm Oskar Tropitzsch GmbH (Marktredwitz, Germany) and ECSI#6 was a kind gift from Dr. Sándor Antus (University of Debrecen, Hungary). [^3^H]5-HT (serotonin, 41.3 Ci/mmol) and [^3^H]citalopram (80 Ci/mmol) were purchased from PerkinElmer Life Sciences (Rodgau, Germany). Scintillation mixture (Rotiszint ecoplus) was purchased from Carl Roth GmbH (Karlsruhe, Germany). Cell culture media were obtained from Sigma Aldrich (St. Louis, MO, USA). The anti-GFP antibody (rabbit, ab290) was from Abcam (Cambridge, UK). An antibody raised against an N-terminal peptide of the G protein β subunit [15] was used to verify comparable loading of lanes. The secondary antibody (Donkey anti-rabbit, IRDye 680RD) was obtained from LI-COR Biotechnology GmbH (Bad Homburg, Germany). All other chemicals were of analytical grade.

### [^3^H]Citalopram binding

Membranes were prepared from HEK293 cells stably expressing GFP-SERT. The cells were washed twice in phosphate-buffered saline (137 mM NaCl, 2.7 mM KCl, 4.3 mM Na_2_HPO_4_, 1.5 mM KH_2_PO_4_, pH adjusted to 7.4), harvested and centrifuged at 2000 rpm for 10 min. The pellets were resuspended in a buffer containing 20 mM HEPES, 2 mM MgCl_2_, 1 mm EDTA, pH adjusted to 7.4 with NaOH, subjected to freeze-thaw cycles in liquid nitrogen and sonicated on ice three times for 10 s with 30 s intervals. Following another centrifugation step, the whole cell membrane pellets were resuspended in buffer and protein concentration was estimated by dye binding (Coomassie Brilliant Blue R-250; Bio-Rad, USA). The binding reaction was carried out in a final volume of 0.1 ml of Na^+^-containing buffer (20 mM Tris-HCl, pH 7.4, 1 mM EDTA, 2 mM MgCl_2_, 120 mM NaCl) containing either 0, 1 or 10 μM cold 5-HT, membranes (2.5 μg/assay), [^3^H]citalopram (3 nM), and the logarithmically spaced concentrations of cocaine (0.1–100 μM), noribogaine (0.1–100 μM) and ECSI#6 (0.1 to 100 μM) at 20 °C for 1 h. Binding reactions were also done in the presence of 120 mM K^+^ buffer (buffer composition: 20 mM Tris-HCl, pH 7.4, 1 mM EDTA, 2 mM MgCl_2_, 120 mM KCl; 4 mM Na^+^ were present from the carry-over of Hepes.NaOH and from Na.EDTA). For binding reactions under these conditions, 7-8 μg of membranes were used to improve the dynamic range of binding. The binding reactions were terminated by harvesting the membranes on glass fiber filters precoated with polyethyleneimine and rapid washing with ice-cold wash buffer (10 mM Tris.HCl, pH 7.4, 120 mM NaCl, 2 mM MgCl2). The radioactivity trapped on the filters was quantified by liquid scintillation counting. Nonspecific binding was measured in the presence of 10 μM paroxetine.

### [^3^H]5-HT uptake

For all uptake assays, HEK293 cells expressing wild-type human YFP-tagged SERT were seeded on poly-D-lysine-coated 96-well plates at a density of 30,000 cells/well. After 24 h, the medium was removed and washed once with Krebs-Hepes buffer (10 mM HEPES.NaOH, pH 7.4, 120 mM NaCl, 3 mM KCl, 2 mM CaCl2, 2 mM MgCl2, and 2 mM glucose). For uptake inhibition assays, logarithmically spaced concentrations of cocaine (0.1–100 μM), noribogaine (0.1–100 μM) or ECSI#6 (0.3–300 μM) were prepared in a buffer containing either 0, 1 or 10 μM cold 5-HT and added to the washed cells for 10 min as a pre-incubation step (50 μl/assay). The same logarithmically spaced concentrations of the 3 compounds were prepared in a buffer containing either 0.2 μM [^3^H]5-HT, or 0.2 μM [^3^H]5-HT with 1 μM cold 5-HT or 0.2 μM [^3^H]5-HT with 10 μM cold 5-HT. After pre-incubation for 10 min, the radioactivity was added for 1 min to achieve a final volume of 100 μl/ well and a final [^3^H]5-HT concentration of 0.1 μM. The reaction was terminated after 1 min by aspiration of the reaction medium followed by a single wash with ice-cold buffer. For the determination of K_M_ and maximal velocity V_m_ax of 5-HT transport by SERT, YFP-SERT cells were pre-incubated for 10 min with 50 μl of either buffer in the absence and presence of either cocaine (3, 10 or 30 μM), noribogaine (1, 3 or 10 μM) or ECSI#6 (10, 30 or 100 μM). After the pre-incubation, 5-HT saturation experiments were undertaken by adjusting the specific activity of 0.1 μM [^3^H]5-HT with unlabeled 5-HT to vary between 50 cpm/fmol to 50 cpm/pmol as follows: logarithmically spaced concentrations of cold 5-HT were prepared in a buffer containing either twice the indicated concentrations of [^3^H]5-HT in the absence and presence of cocaine (3, 10 or 30 μM), noribogaine (1, 3 or 10 μM) or ECSI#6 (10, 30 or 100 μM). These solutions (50 μl) were added for 1 min to achieve a final volume of 100 μl/ well and a final [^3^H]5-HT concentration of 0.1 μM. The reaction was terminated after 1 min by aspiration of the reaction medium followed by a single wash with ice-cold buffer.

For uptake assays determining the functional rescue of mutant transporter, HEK293 cells were transfected with either YFP-tagged SERT-PG^601,602^ AA or YFP-tagged WT-SERT plasmids. Transfected cells were seeded on poly-D-lysine-coated 96-well plates at a density of ~60-80,000 cells/well in either the absence or presence of increasing concentrations (0.3-100 μM) of either cocaine, noribogaine or ECSI#6. After 24 h, the cells were washed four times with Krebs-MES buffer (10 mM 2-(N-morpholino)ethanesulfonic acid, pH 5.5, 120 mM NaCl, 3 mM KCl, 2 mM CaCl2, 2 mM MgCl2, and 2 mM glucose) in a 10 min interval and once with Krebs-HEPES (pH 7.4) buffer. The cells were subsequently incubated with 0.2 μM of [^3^H]5-HT for 1 min and the uptake assay was carried out as outlined above. Nonspecific uptake for all uptake experiments was defined in the presence of 10 μM paroxetine. After the uptake reaction, the cells were then lysed with 1% SDS to release the retained radioactivity, which was quantified by liquid scintillation counting.

### Immunoblotting

HEK293 cells were transiently transfected with plasmids encoding either WT-YFP-SERT or YFP-SERT-PG^601,602^AA. Approximately 1.5-2 × 10^6^ of these transfected cells were seeded in 6- well plates in the presence of either cocaine, noribogaine or ECSI#6 (0.3-30 μM). After 24 h, cells were washed thrice with ice-cold phosphate-buffered saline, detached by mechanical scraping, and harvested by centrifugation at 1000 g for 5 min. The cell pellet was lysed in a buffer containing Tris·HCl, pH 8.0, 150 mm NaCl, 1% dodecyl maltoside, 1 mm EDTA, and protease inhibitors (Complete, Roche Applied Science). This soluble protein lysate was separated from the detergent-insoluble material by centrifugation (16,000g for 15 min at 4 °C). An aliquot of this lysate (20 μg) was mixed with sample buffer containing 1% SDS and 20 mM DTT, denatured at 45 °C for 30 min and resolved in denaturing polyacrylamide gels. After protein transfer onto nitrocellulose membranes, the blots were probed with an antibody against GFP (rabbit, ab290) at a 1:3000 dilution overnight. This immunoreactivity was detected by fluorescence detection using a donkey anti-rabbit secondary antibody at 1:5000 dilution (IRDye 680RD, LICOR). The lower part of the blot was also probed with the antibody recognizing the G protein β subunits to verify equal loading.

### Whole-cell patch clamp recordings

HEK293 cells stably expressing wild-type GFP-tagged human SERT were seeded at low density on poly-D-lysine coated dishes. 24h after seeding, these cells were subjected to patch clamp recordings in the whole cell configuration. In most instances cells were continuously maintained in an external solution containing 140 mM NaCl, 3 mM KCl, 2.5 mM CaCl_2_, 2 mM MgCl_2_, 20 mM glucose, and 10 mM HEPES (pH adjusted to 7.4 with NaOH) and the drugs were diluted therein. The internal solution in the patch pipette contained 133 mM potassium gluconate (CH_2_OH(CHOH)_4_COOK), 5.9 mM NaCl, 1 mM CaCl_2_, 0.7 mM MgCl_2_, 10 mM HEPES, 10 mM EGTA (pH adjusted to 7.2 with KOH). Drugs were applied using a 4-tube ALA perfusion manifold (NPI Electronic GmbH, Germany) and a DAD-12 superfusion system (Adams & List, Westbury, NY) allowing for complete solution exchange around the cells within 100 ms. Current amplitudes and associated kinetics were quantified using Clampfit 10.2 software. Passive holding currents were subtracted, and the traces were filtered using a 100-Hz digital Gaussian low-pass filter.

### Drosophila genetics and drug treatment

The transgenic UAS reporter lines for YFP-tagged human wild-type SERT and SERT-PG^601,602^AA were generated using the pUAST-attB vector (gift from Drs. Bischof and Basler, University of Zürich) and injected into embryos of ZH-86Fb flies (Bloomington stock no. 24749). Positive transformants were isolated and selected. TRH-T2A-Gal4 (Bloomington stock no. 84694) was used to drive the expression of transporters in serotonergic neurons. Three-day old male; TRH-T2A-Gal4/UAS-YFP-hSERT-WT; or TRH-T2A-Gal4/UAS-YFP-hSERT-PG^601,602^ AA; flies were treated with food supplemented with 100 μM noribogaine or 100 μM ECSI#6 for 48 h. The brains of these flies were then imaged using confocal microscopy. All flies were kept at 25 °C, and all crosses were performed at 25 °C.

### Immunohistochemistry and imaging

Adult fly brains were dissected in phosphate-buffered saline (PBS) and fixed in 2% paraformaldehyde in PBS for 1 h at room temperature. Brains were then washed three times in 0.1% Triton X-100 in PBS for 20 min on a shaker. Blocking was performed in 10% goat serum for 1 h at room temperature on a shaker. Brains were then incubated in primary antibody overnight in PBS containing 3% BSA and 0.3% Triton X-100 at 4 °C on a shaker. The rabbit polyclonal IgG directed against GFP (1:1000 dilution; A-11122, Invitrogen) and anti-serotonin antibody (S5545, Sigma Aldrich) were used as primary antibodies. After three washes for 20 min with PBS containing 0.1% Triton X-100, the brains were incubated overnight at 4 °C with a secondary antibody in PBS containing 0.3% Triton X-100 on a shaker. Alexa Fluor 488- or 568-labeled goat anti-rabbit IgG (1: 500 or 1:300 Invitrogen) was used as a secondary antibody. Following incubation with secondary antibody, the brains were washed three times with PBS containing 0.1% Triton X-100 and were mounted using Vectashield® (Vector Laboratory, Burlingame, CA). Images were captured on a Leica SP5II confocal microscope with 20-fold magnification. Z-stack images were scanned at 1.5-μm section intervals with a resolution of 512 × 512 pixels. Images were processed with ImageJ.

### Kinetic modelling

A kinetic model of the transport cycle of SERT was built based on the reaction scheme in Fig.7A. This scheme is simplified because (i) it ignores Cl^-^ and proton binding to SERT, (ii) it does not account for voltage dependence and (iii) it assumes a sequential-instead of a random binding order of Na^+^ and substrate. These simplifications were justified by the fact that in the experiments, which we simulated we did not change the Cl-/proton concentrations in the bath or the electrode solution, we did not apply voltage jumps and we used saturating concentrations of Na^+^ throughout. The time-dependent changes in state occupancies were evaluated by numerical integration of the resulting system of differential equations using the Systems Biology Toolbox [16] in Matlab 2019b (The MathWorks, Inc., Natick, Massachusetts). We used this model to evaluate how state-dependent binding of an inhibitor impinges on substrate uptake. To this end, we allowed for the inhibitor to bind to each individual state specified in the reaction scheme at a time. The rate of substrate uptake was modeled as: TiS*koff-Ti*kon*Sin; where TiS and Ti are the occupancies of the substrate-bound and apo inward facing conformation, respectively and k_off_ and k_on_ the dissociation and association rate of substrate and S_in_ the intracellular substrate concentration. The numeric values used to parameterize the microscopic rate constants of the model can be found in the legend of Fig.7A.

## Results

### Cocaine, noribogaine and ECSI#6 exhibit differences in the ability to block SERT uptake

Affinity estimates of exogenous ligands to monoamine transporters can be evaluated by their ability to block either the functional uptake of a tritiated substrate or binding of a high affinity radiolabeled inhibitor to the transporter. For the uptake inhibition experiments, we calculated IC_50s_ for the ability of cocaine, noribogaine or ECSI#6 to block transport of 0.1 μM [^3^H]5-HT by SERT either in the absence (IC_50(0)_) or prior pre-incubation of either 1 μM (IC_50(1)_) or 10 μM (IC_50(10)_) cold 5-HT. Cocaine blocks SERT uptake of [^3^H]5-HT with an IC_50(0)_ of 9.96 μM (95% CI: 8.7 to 11.4, Fig.1A, solid blue curve). The IC_50(1)_ was indistinguishable from IC_50(0)_ (Fig.1A, blue dashed curve, IC_50_ = 9.8 μM, 95% CI: 7.1 to 13.6), while IC_50(10)_ was right shifted by ~ 4-fold (Fig.1A, blue dotted curve, IC_50_ = 34.6 μM, 95% CI: 24.8 to 48.3). This right shift is expected and readily explained; increasing concentrations of unlabeled 5-HT in the uptake inhibition assay inhibits cocaine binding to SERT in a competitive manner. Noribogaine, on the other hand, blocked substrate uptake of SERT more potently in the presence of unlabeled 5-HT (*cf.* solid, dashed and dotted green lines, Fig.1B, IC_50(0)_: 5.68 μM [95% CI: 4.36 to 7.39], IC_50(1)_: 2.39 μM [95% CI: 1.84 to 3.09], IC_50(10)_: 1.89 μM [95% CI: 1.46 to 2.44]). In the presence of unlabeled 5-HT, this left shift in potency was more prominent with ECSI#6 (*cf.* solid, dashed and dotted red lines, Fig. 1B, IC_50(0)_: 440.5 μM [95% CI: 325.8 to 595.6], IC_50(1)_: 74.14 μM [95% CI: 55.4 to 99.22], IC_50(10)_: 19.28 μM [95% CI: 13.23 to 28.09]). These observations clearly show that noribogaine and ECSI#6 bind to SERT with higher affinity in the presence of a substrate. Thus, their mode of binding to the transporter differs from that of cocaine.

**Fig 1.**
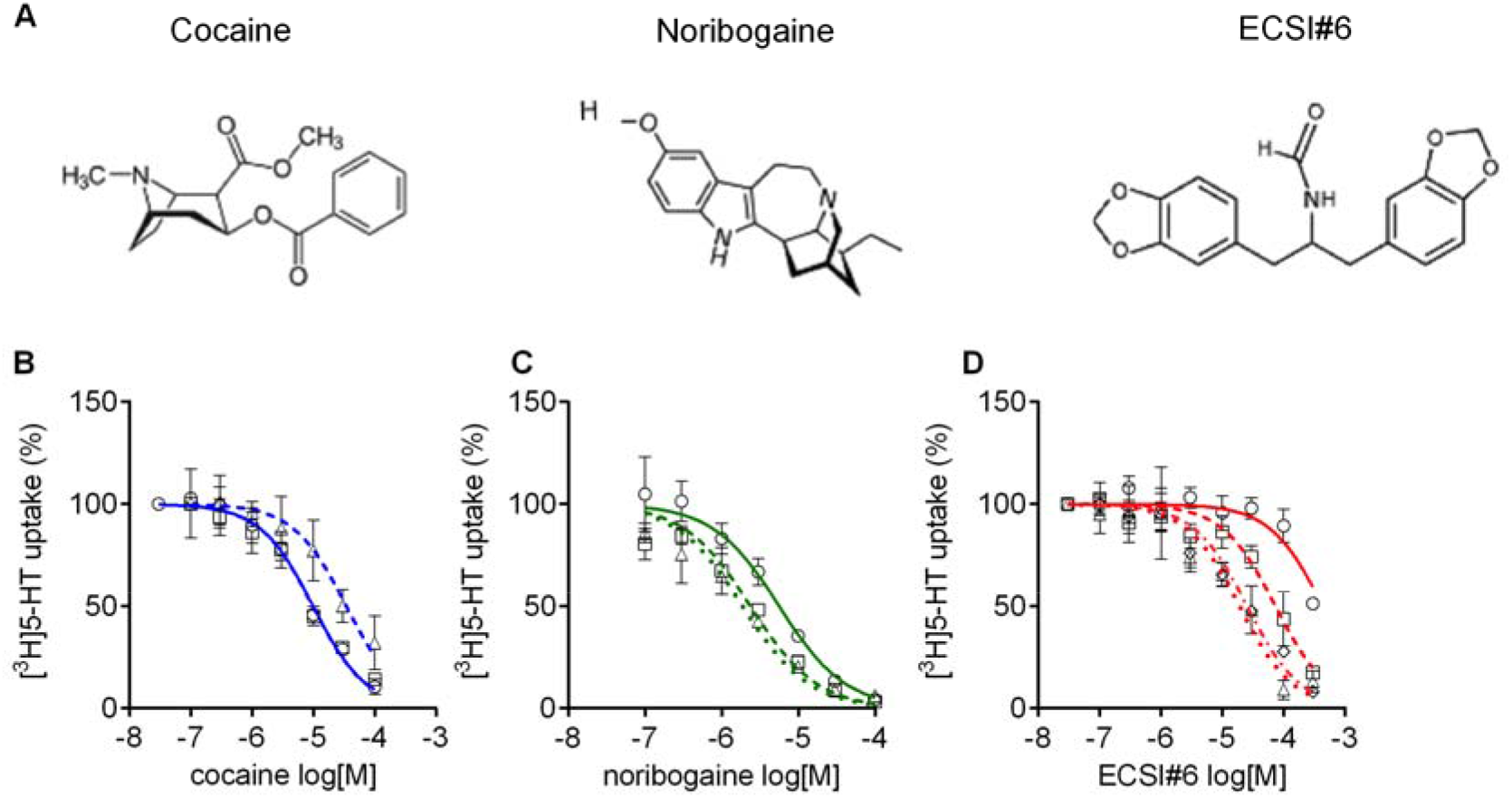
Inhibition of SERT mediated [3H]5-HT uptake by cocaine, noribogaine and ECSI#6. **A**, the chemical structures of cocaine, noribogaine and ECSI#6. **B - D**, HEK293 cells stably expressing wild-type YFP-SERT (~30,000 per well) were seeded onto 96-well plates. After 24 hours, inhibition of substrate uptake by cocaine (B), noribogaine (C) or ECSI#6 (D) was determined as outlined under *Materials and Methods* in the absence of additional unlabeled 5-HT (circles & solid lines, IC_50(0.1)_ or in the presence of 1 μM (squares and dashed lines, IC_50(1)_), 3 μM (diamonds and dashed-dotted lines in D; IC_50(3)_), or 10 μM 5-HT (triangles, dotted lines, IC_50(10)_) of unlabeled 5-HT. Paroxetine (10 μM) was used to determine nonspecific uptake, which was ~5% of total uptake. Uptake was normalized to the specific uptake in the absence of inhibitors (4.2 ± 0.6 pmol.min^-1^.10^-6^ cells), which was set to 100% (i.e., no inhibition) to account for inter-experimental variations. Data represented are the means ± S.D. (error bars) from three independent experiments done in triplicate. The curves were generated by fitting a sigmoidal function through data points normalized between 100% and no uptake. The concentrations giving half maximum inhibition (IC_50_)were (means and 95% confidence interval in parenthesis): cocaine - IC_50(0.1)_ = 9.9 μM (8.7 – 11.4), IC_50(1)_ = 9.8 μM (7.1 – 13.5), IC_50(10)_ = 34.6 μM (24.8 – 48.3); noribogaine - IC_50(0.1)_ = 5.7 μM (4.3 – 7.3), IC_50(1)_ = 2.4 μM (1.8 – 3.1), IC_50(10)_ = 1.9 μM (1.5 – 2.4); ECSI#6 - IC_50(0.1)_ = 440.5 μM (325.8 – 595.6), IC_50(1)_ = 74.1 μM (55.4 – 99.2), IC_50(3)_ = 25.6 μM (19.1 – 34.2), IC_50(10)_ = 19.3 μM (13.2 – 28.1).

### Cocaine, noribogaine and ECSI#6 bind to different SERT conformational intermediates

The data summarized in Fig. 1 indicate that ECSI#6 and noribogaine differ in their mode of binding from cocaine. There are two explanations for the left shift in the uptake inhibition by by ECSI#6 (and by noribogaine) in the presence of increasing concentrations of 5-HT: (i) binding of 5-HT to SERT results in a conformational transition, which exposes a binding pocket for ECSI#6 (and for noribogaine). (ii) Alternatively, uptake of 5-HT by SERT results in the accumulation of the K^+^-bound conformational intermediate(s), which display a higher affinity for ECSI#6 or noribogaine. The first model posits that serotonin and ECSI#6 are bound simultaneously. This has been observed with serotonin and carbamazepine, which can bind concomitantly to the outward-facing state of SERT [17]. To address which of the two possibilities can explain the experimental observations, membranes harboring SERT were incubated with 3 nM [^3^H] citalopram and logarithmically spaced concentrations of ECSI#6 (and the reference compounds cocaine and noribogaine) in the presence of 120 mM NaCl, which promotes the outward-facing conformation. It is evident from Fig. 2C that addition of 1 or 10 μM 5-HT shifted the displacement curve of ECSI#6 to the right to an extent which was comparable to that seen with cocaine (Fig. 2A) and noribogaine (Fig. 2B). Thus, binding to SERT of serotonin and of these three ligands was mutually exclusive.

**Fig 2.**
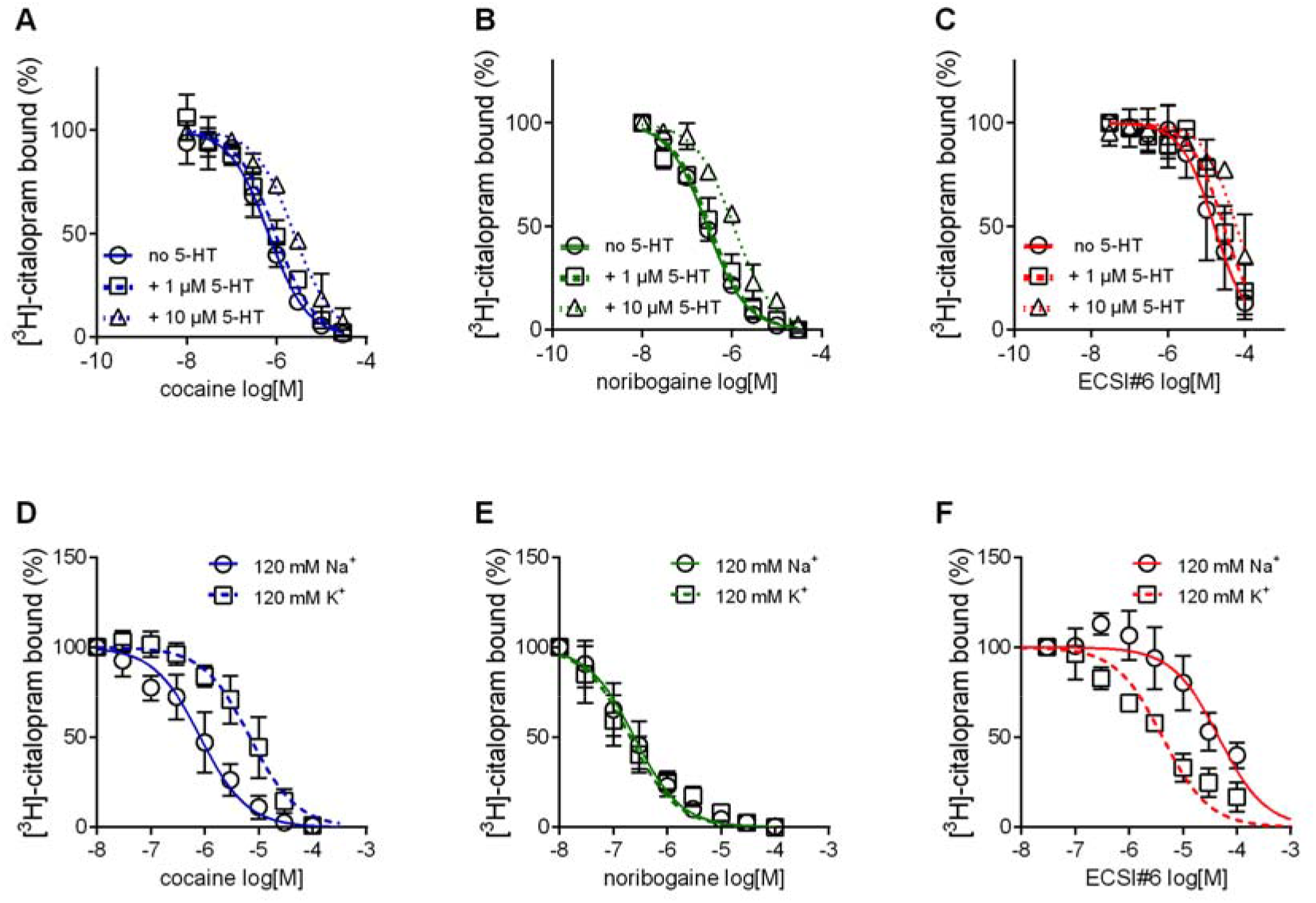
Inhibition of [^3^H]citalopram binding by cocaine, noribogaine and ECSI#6. **A – G,** Inhibition of [^3^H]Citalopram binding to SERT by cocaine (A, D; blue curves), noribogaine (B, E; green curves) or ECSI#6 (C, F; red curves). The binding inhibition assays in **A-C** were performed in the absence (open circles, solid lines) and presence of 1 μM (squares, dashed lines) or of 10 μM 5-HT (triangles, dotted lines) in buffer containing 120 mM Na^+^ with 2-3 μg membranes (0 and 1 μM 5-HT) or 7-8 μg membranes (10 μM 5-HT). The specific binding values for assays in the absence (no 5-HT) or presence of 1 μM and 10 μM 5-HT were 28 ± 5, 17 ± 3 and 21 ± 0.3 fmol/assay, respectively. These values were normalized to 100% to account for inter-experiment variability. The binding inhibition assays in **D-F** were performed in a binding buffer that contained either 120 mM Na^+^ (circles, solid curves) or 120 mM K^+^ (squares, dashed curves). Assays in the presence of 120 mM NaCl were done with 2-3 μg membranes; for assays with 120 mM KCl, 7-8 μg of membranes were used to improve dynamic binding range. The specific binding values for assays with the 120 mM NaCl and 120 mM KCl buffer were 25 ± 14 and 21 ±5 fmol/assay, respectively. These values were normalized to 100% to account for inter-experiment variability. All data are the means from 3 independent experiments done in duplicate; error bars represent S.D. The curves generated were generated by fitting the data points to the equation for a monophasic inhibition. The concentrations giving half maximum inhibition (IC_50_ values) were calculated from each plot yielding (95% confidence intervals in parenthesis): **A,** cocaine - IC_50(no 5-HT)_ = 0.66 μM (0.54 – 0.82), IC_50(1 μM 5-HT)_ = 0.94 μM (0.76 – 1.09), IC_50(10 μM 5-HT)_ = 2.48 μM (2.08 – 2.90); **B,** noribogaine - IC_50(no 5-HT)_ = 0.28 μM (0.24 – 0.31), IC_50(1 μM 5-HT)_ = 0.32 μM (0.26 – 0.42), IC_50(10 μM 5-HT)_ = 1.10 μM (0.92 – 1.31); **C,** ECSI#6 - IC_50(no 5-HT)_ = 16.02 μM (10.96 – 22.42), IC_50(1 μM 5-HT)_ = 28.6 μM (20.2 – 40.0), IC_50(10 μM 5-HT)_ = 60.0 μM (38.2 – 94.2); **D,** cocaine – IC_50 (120 Na^+^)_ = 0.82 μM (0.62 – 1.10), IC_50(120 K^+^)_ = 7.94 μM (5.34 – 9.01); **E,** noribogaine – IC_50 (120 Na^+^)_ = 0.24 μM (0.18 – 0.32), IC_50(120 K^+^)_ = 0.20 μM (0.16 – 0.24); **F,** ECSI#6 - IC_50(120 Na^+^)_ = 46.2 μM (31.6 – 67.6), IC_50(120 K^+^)_ = 4.18 μM (2.58 – 5.88). Paroxetine (10 μM) was used to determine nonspecific binding, which was ≤10% of total binding.

We also explored the alternative hypothesis, i.e. that ECSI#6 bound preferentially to a K^+^-bound state of SERT. K^+^ promotes the return step of SERT [2,18]. Accordingly, in the presence of 120 mM K^+^, appreciable binding of radioligand can be measured [19]. We compared the ability of ECSI#6 and of the reference compounds cocaine and noribogaine to displace [^3^H] citalopram in the presence of 120 mM NaCl and 120 mM KCl. As expected, cocaine was substantially less potent (by an order of magnitude) in inhibiting [^3^H]citalopram binding in the presence of 120 mM KCl than in the presence of 120 mM NaCl (Fig. 2D). This indicates that cocaine binds more readily to Na^+^-bound SERT. This distinct binding preference was not obvious with noribogaine (Fig. 2E). Importantly and in contrast to cocaine, ECSI#6 was about 10-fold more potent in displacing [^3^H] citalopram binding in the presence of 120 mM KCl than in the presence of 120 mM NaCl (Fig. 2E). Thus ECSI#6 clearly preferred K^+^-bound SERT.

### Cocaine, noribogaine and ECSI#6 differ in their mode of SERT inhibition

Cocaine and noribogaine are competitive and non-competitive inhibitors of substrate uptake by SERT, respectively [20,21]. The mode of inhibition by ECSI#6 was anticipated to differ from either compound because of the pronounced left-shift in both, the uptake inhibition curve of ECSI#6 in the presence of increasing serotonin (Fig. 1D) and the radioligand displacement curve in the presence of high K^+^ (Fig. 2F). This was the case: increasing concentrations of ECSI#6, shifted the saturation hyperbolae for 5-HT uptake (Fig. 3C, top) and caused a reduction in both, K_M_ (Fig. 3C middle panel) and V_MAX_ (Fig. 3C, bottom panel). This is the hallmark of uncompetitive inhibition. In parallel experiments, we recapitulated competitive and non-competitive inhibition of substrate uptake by cocaine (Fig. 3A) and by noribogaine (Fig. 3B), respectively.

**Fig 3.**
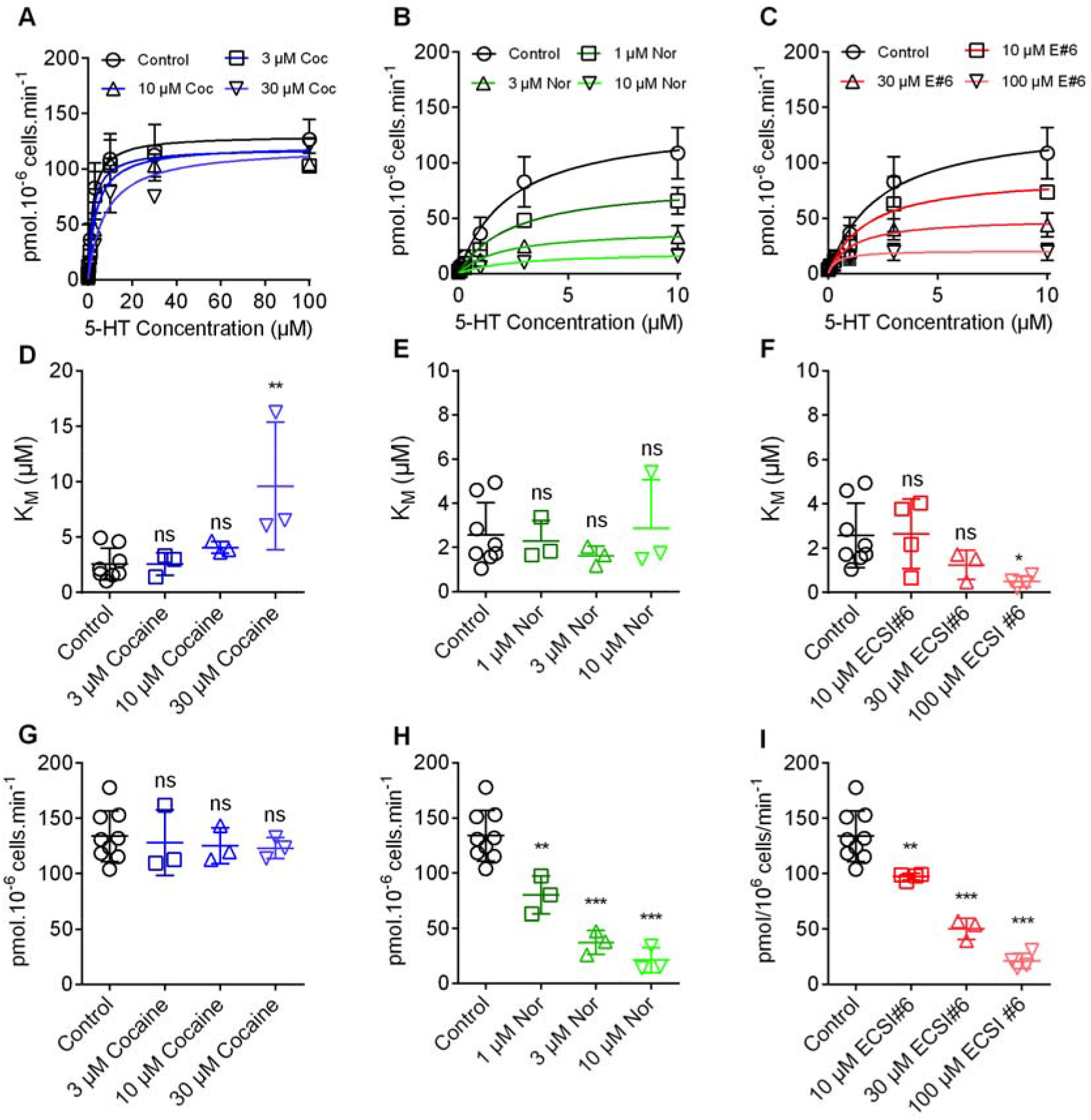
Different modes of inhibition by cocaine, noribogaine and ECSI#6 of substrate uptake by SERT. HEK293 cells stably expressing YFP-SERT (30,000/well) were pre-incubated in buffer in the absence (control, black circles) or presence of the indicated concentrations of cocaine (A), noribogaine (B) or ECSI#6 (C) for 10 min; subsequently, the uptake reaction was initiated as outlined in Materials and Methods. In the absence of any inhibitor (black circles and black lines, control), the K_M_ and V_MAX_ of 5-HT transport by SERT was 2.3 μM (95% CI, 1.5 – 3.2) and 137.3 pmol.min^-1^.10^-6^ cells (95% CI, 117.4 – 157.3); these control curves are identical in all panels. **A,** In the presence of 3 μM (blue squares), 10 μM (blue upward triangles) and 30 μM cocaine (blue downward triangles) there was a progressive increase in K_M_-values (panel D with K_M_ = 2.2 μM [95% CI, 1.1 – 3.3], 3.7 μM [95% CI, 1.9 – 5.5] and 8.1 μM [95% CI, 1.9 – 5.5], respectively), but V_max_-values remained constant (panel G, V_MAX_ = 118.3 pmol.min^-1^.10^-6^ cells [95% CI, 117.4 – 157.3], 121.1 pmol.min^-1^.10^-6^ cells [95% CI, 105.5 – 136.8]) and 120.1 pmol.min^-1^.10^-6^ cells [95% CI, 104.6 – 135.6], respectively). **B,** Preincubation with 1 μM (green squares), 3 μM (green upward triangles) and 10 μM noribogaine (green downward triangles) did not change the K_M_-values (panel E with K_M_ = 2.4 μM [95% CI, 1.4 – 3.5], K_M_ = 1.9 μM [95% CI, 0.6 – 3.3] and 2.4 μM [95% CI, 0.6 – 5.5], respectively), but resulted in a concentration-dependent reduction in V_MAX_ (panel H with V_MAX_ = 82.4 pmol.min^-1^.10^-6^ cells [95% CI, 69.3 – 95.4]), 39.9 pmol.min^-1^.10^-6^ cells [95% CI, 30.6 – 49.2]) and 19.5 pmol.min^-1^.10^-6^ cells [95% CI, 13.9 – 24.9], respectively). **C,** Preincubation with 10 μM (red squares), 30 μM (red upward triangles) and 100 μM ECSI#6 (red downward triangles) led to a drop in both, the K_M_-values (panel F; K_M_ = 1.7 μM [95% CI, 0.5 – 2.9], 1.1 μM [95% CI, 0.2 – 1.9] and 0.4 μM [95% CI, 0.0 – 0.8], respectively) and the V_MAX_-values (panel H, V_MAX_ = 89.5 pmol.min^-1^.10^-6^ cells [95% CI, 68.5 – 110.0]), 49.9 pmol.min^-1^.10^-6^ cells [95% CI, 37.5 – 62.3] and 20.8 pmol.min^-1^.10^-6^ cells [95% CI, 15.6 – 26.0], respectively). Data are the means + S.D. (error bars) from at least 3 independent experiments done in triplicate. The curves were generated by fitting the data points to the equation for a rectangular hyperbola. Control K_M_- and V_MAX_-values were pooled and are hence the same in panels D-F and G-I, respectively. Statistical comparisons were done by one-way ANOVA followed by Dunnett’s post-hoc test to verify significant differences vs control (*p<0.05, **p<0.01, ***p<0.001).

### Kinetics of current inhibition by ECSI#6

The different mode of inhibition, which was observed for ECSI#6, implies that it affects one or several partial reactions of the transport cycle in manner distinct from cocaine and noribogaine. The transport cycle of SERT can be addressed by whole-cell patch-clamp recordings; electrophysiological recordings of currents through SERT provide unmatched temporal resolution of ligand and co-substrate binding and unbinding events [18,22]. When challenged with a substrate like 5-HT or amphetamines, SERT carries inward currents that are comprised of an initial transient peak current and a steady current, which reflect the initial synchronized movement of substrate and co-substrate through the membrane electric field and the continuous cycling of the transporter in the forward transport mode, respectively [23,24]. In contrast, inhibitors only elicit a minor peak current on application to SERT expressing cells [25]. As shown in Fig. 4A, cocaine and noribogaine induced transient peak currents, while ECSI#6 failed to do so. We relied on measuring the time course of inhibition of the substrate-induced steady current through SERT to infer the kinetics of inhibitor binding: we first applied 5-HT at a saturating concentration, which elicited the peak current and the steady state current (Fig. 4B). After 5 seconds, the cells were superfused with inhibitors in the continuous presence of the substrate. ECSI#6 and the reference compounds cocaine and noribogaine (red, blue and green trace, respectively in the magnified section in Fig. 4B) led to a rapid inhibition of the substrate-induced current, which was rapidly reversed by removal of the inhibitor. The inhibitor-induced suppression of the steady current was adequately described by a monoexponential decay. The rate constant reflects the apparent rate of association (k_app_) of the inhibitors. In a simple bimolecular reaction (corresponding to binding of the inhibitor to SERT), raising the inhibitor concentration is predicted to result in accelerated binding. This was not the case for cocaine (Fig. 4C): as shown previously [21], the k_app_ of cocaine only increased in the very low concentration range and leveled off at about 2 s^-1^. This is expected for a typical inhibitor, which interacts with the outward-facing conformation of SERT, because binding is limited by the return step from the inward-facing to the outward-facing state. This is the rate-limiting reaction in the transport cycle of SERT [22,23]. In contrast, k_app_ for both, noribogaine (Fig. 4D) and ECSI#6 (Fig.4E), increased in a linear manner as their concentrations were raised. This can only be achieved, if noribogaine and ECSI#6 bind to the inward-facing conformation of SERT directly. Noribogaine was ~10-fold more potent than ECSI#6 in imposing the block on the substrate-induced current through SERT.

**Fig 4.**
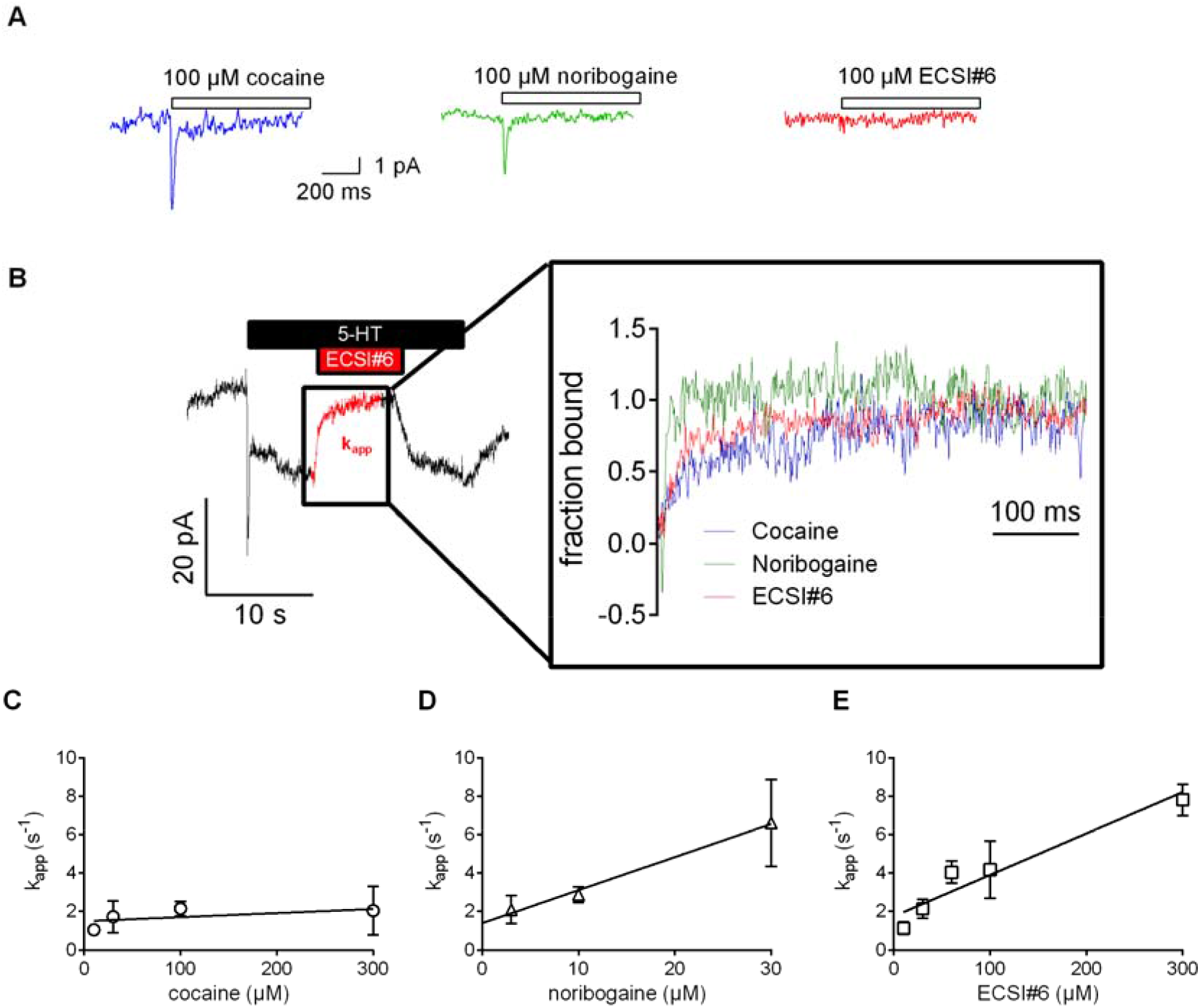
Kinetics of current inhibition by cocaine, noribogaine and ECSI#6. Single HEK293 cells stably expressing GFP-SERT were voltage-clamped to −60 mV using the whole cell patch clamp technique under physiological ionic gradients. **A,** representative recording of a cell superfused with cocaine (100 μM, left-hand trace), noribogaine (100 μM, middle trace) or ECSI#6 (100 μM, righthand trace). **B,** Cells were initially challenged with 10 μM 5-HT. After 5 seconds, the cells were superfused with varying concentrations of either cocaine, noribogaine or ECSI#6 for 5 seconds in the continuous presence of 10 μM 5-HT, which resulted in current inhibition. Thereafter, the compounds were removed by superfusion with 10 μM 5-HT alone for 10 s to monitor recovery of the steady current. The left-hand panel shows the original trace of a representative current recorded after sequential superfusion with 10 μM 5-HT for 5 s, which triggered a peak current followed by a steady current, 100 μM ECSI#6 in the continuous presence of 10 μM 5-HT, which led to complete suppression of the current (highlighted in red), and 10 μM 5-HT, which resulted in recovery of the steady current. The right-hand panel shows magnified representative segments of the traces recorded from different cells, which had been superfused with either cocaine (blue, 100 μM), noribogaine (green, 30 μM) or ECSI#6 (red, 100 μM). The time course of current inhibition was fitted to the equation for a mono-exponential decay to estimate the apparent rate constant (k_app_). **C, D, E**; Summary of the analysis of the kinetics of current inhibition by cocaine (open circles), noribogaine (open triangles) and ECSI#6 (open squares), respectively. The k_app_-values were derived from experiments done as in panel A with the indicated concentrations of the compounds. Data are means ± S.D. ≥5 independent recordings. The k_app_-values were plotted against the concentration. The slopes of the resulting lines were calculated by linear regression. The slope for cocaine did not deviate in a statistically significant manner from zero (F-test; p = 0.12), while those for noribogaine and ECSI#6 did (p<0.0001 in both instances).

### ECSI#6 can rescue a folding-deficient SERT mutant

In the endoplasmic reticulum (ER), the folding trajectory of SERT moves through the inward-facing conformation [26,27]. Compounds, which bind to the inward-facing conformation, act as pharmacochaperones: they rescue folding-deficient mutants of SERT and of DAT [11]. Taken together, the experiments summarized in Figs. 1 to 4 highlight that ECSI#6 binds to SERT in a unique mode, which favors the K^+^-bound inward-facing state. Thus, ECSI#6 was predicted to have pharmacochaperoning activity. This prediction was verified by using SERT-PG^601,602^AA. This mutant has a severe folding defect, the bulk of the protein is trapped in the ER at an early stage of the SERT folding trajectory [28,29]. Accordingly, only a small fraction of SERT-PG^601,602^AA reaches the cell surface, where it supports substrate uptake, but the folding defect can be corrected, if cells are pre-incubated with noribogaine [29]: as shown in Fig. 5A, in paired transient transfections, in HEK293 cells expressing SERT-PG^601,602^AA, substrate uptake was only ~10% of cells expressing wild type SERT. If HEK293 cells were preincubated with 30 μM of cocaine, noribogaine or ECSI#6, appreciable functional rescue was observed with ECSI#6 and the reference compound noribogaine; cocaine was ineffective (Fig. 5B). This pharmacochaperoning action of ECSI#6 and of noribogaine was concentration-dependent and saturable (Fig. 5C): noribogaine was more potent (EC_50_ = 0.46 μM) and more efficacious (E_max_= 69.5% of the transport velocity observed in cells expressing wild type SERT) than ECSI#6 (EC_50_ = 2.9 μM; E_max_= 52%).

**Fig 5.**
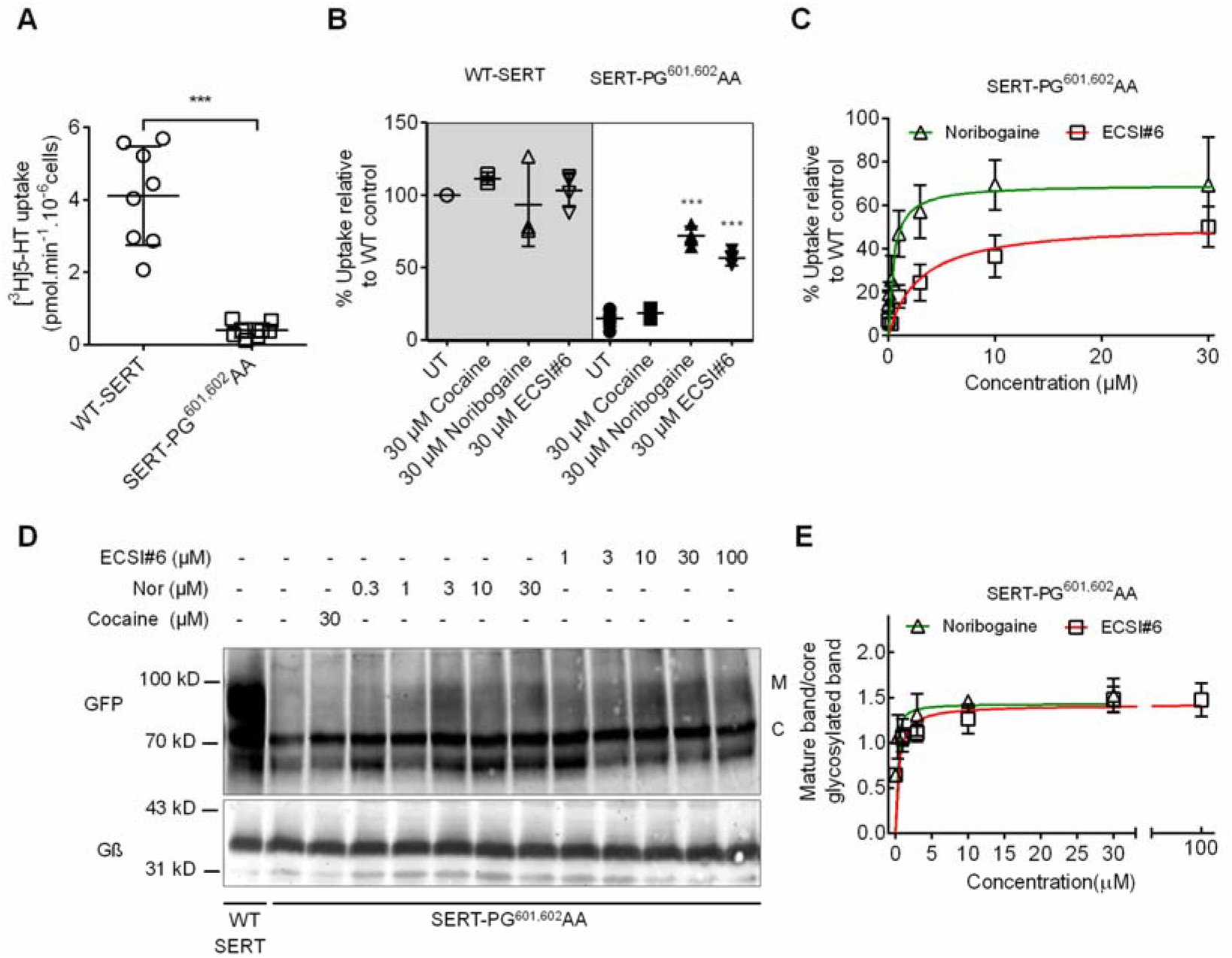
[^3^H]5-HT uptake by and glycosylation pattern of hSERT-PG^601,602^AA expressed in HEK293 cells incubated with cocaine, noribogaine or ECSI#6. **A**, Comparison of [^3^H]5-HT uptake by HEK293 cells transiently expressing either wild-type SERT (WT SERT) or SERT-PG^601,602^AA. Cellular uptake of substrate was measured with 0.1 μM[^3^H]5-HT as outlined under Materials and Methods and amounted to (means ± SD) 4.1 + 1.3 pmol.min^-1^.10^-6^ cells and 0.39 + 0.20 pmol.min^-1^.10^-6^ cells for wild-type SERT and SERT-PG^601,602^ AA, respectively. Each symbol represents the result of an individual experiment (done in triplicate). The statistical comparison was done by a Mann-Whitney test (p = 0.0009). **B**, Cells transiently expressing either WT-SERT or SERT-PG^601,602^AA were incubated in the presence of 30 μM of either cocaine, noribogaine or ECSI#6. After 24 hours, cellular substrate uptake was determined with 0.1 μM [^3^H]5-HT. Uptake values from individual conditions were normalized to the ones from untreated cells expressing wild-type SERT (set to 100%) to account for inter-experimental variations. Values from individual experiments (done in triplicate), represented collectively as a box plot, are shown as mean + S.D as follows: WT-SERT untreated (4.1 + 1.3 pmol.min^-1^.10^-6^ cells set to 100%), WT-SERT + 30 μM cocaine (111.4 + 4.2%), WT-SERT + 30 μM Noribogaine (93.5 + 28.8%), WT-SERT + 30 μM ECSI#6 (103.3 + 11.3%), SERT-PG^601,602^AA untreated (14.8 + 5.3%), SERT-PG^601,602^ AA + 30 μM cocaine (18.6 + 3.9%), SERT-PG^601,602^AA + 30 μM Noribogaine (71.9 + 6.4%), SERT-PG^601^,^602^AA + 30 μM ECSI#6 (56.5 + 5.1%). The statistical comparison of untreated cells expressing SERT-PG^601,602^AA and their treated counterparts was done by one-way ANOVA followed by *post hoc* Dunnett’s multiple comparisons (***, p < 0.001). **C**, Concentration-response curves for pharmacochaperoning of SERT-PG^601,602^AA by noribogaine and ECSI#6. Rescued uptake was normalized to uptake velocity measured in parallel in HEK293 cells transiently expressing WT-SERT to account for inter-experimental variations. E_MAX_ and EC_50_ for noribogaine and ECSI#6 were determined by fitting the data to the equation for a rectangular hyperbola (95% CI in parenthesis): for noribogaine and ECSI#6, E_MAX_ was 69.5% (59.2 – 79.7) and 52.0% (44.6 – 59.5), and EC_50_ was 0.46 μM (0.15 – 0.77) and 2.9 μM (1.1 – 4.7), respectively. The data were obtained in at least three independent experiments carried out in triplicate. The error bars indicate S.D. **D**, Confluent cultures of HEK293 cells transiently expressing SERT-PG^601,602^AA (1 well of a 6 well plate/condition) were treated with either cocaine (30 μM), noribogaine or ECSI#6 in the indicated concentrations for 24 h. Untreated cells were taken as negative controls (second lane in the representative blot). Membrane proteins extracted from these cells were denatured and resolved with SDS-PAGE and transferred onto nitrocellulose membranes. The blots were incubated overnight at 4 °C with anti-GFP (top) or anti-Gβ (bottom, loading control) antibodies. The immunoreactive bands were detected with fluorescently labeled secondary antibodies. The blot is representative of three independent experiments. **E**, The intensities of the immunoreactive bands were quantified by densitometry; the ratio of mature (M) to core glycosylated band (C) was corrected for the intensity of the loading control (Gβ). These normalized values (expressed as A.U. – arbitrary units) were plotted as a function of drug concentration, and fitted to an equation for a rectangular hyperbola. The E_MAX_ and EC_50_ (95% CI in parenthesis) were E_MAX_ = 1.43 A.U. (1.18 – 1.68) and 1.42 (1.18 – 1.65), EC_50_ = 0.14 μM (0.01 – 0.35) and 0.44 μM (0.01 – 1.05) for noribogaines and ECSI#6, respectively.

We independently confirmed that the rescue of SERT-PG^601,602^AA by ECSI#6 (and the reference compound noribogaine) was indeed due to increased ER export of the mutant by determining the glycosylation state of the protein. Membrane proteins acquire N-linked core glycans co-translationally in the ER; in the Golgi apparatus, they incorporate additional sugar moieties to achieve mature glycosylation. Two species were visualized by immunoblotting of whole cell lysates from cells transiently expressing wild-type SERT (lane 1 in representative blot displayed in Fig. 5D): (i) the band at approximately 75 kDa corresponds to the ER-resident core-glycosylated species [28,30]; (ii) a broad smear migrating in the range of 90 to 110 kDa represent the transporters carrying mature glycans [28]. Lysates from cells expressing SERT-PG^601,602^AA alone show the predominant presence of the core glycosylated species indicating ER retention (lane 2, Fig. 5D). If lysates from cells expressing SERT-PG^601,602^AA were treated for 24 hours with noribogaine (lanes 4-8 in Fig. 5D) or with ECSI#6 (lanes 9-13 in Fig. 5D), there was a concentration-dependent increase in the appearance of the mature glycosylated species. Noribogaine was more potent than ECSI#6 (Fig. 5E). We note that the EC_50_-values for enhancing the accumulation of the mature glycosylated SERT (Fig. 5E) differed from those for restoring uptake. This presumably reflects the modest precision from which band densities can be quantified with. As expected, pre-treatment of cells with a saturating concentration of cocaine did not cause any appreciable increase in the SERT species carrying mature glycans (lane 3, Fig. 5D).

Human SERT and DAT can substitute for their orthologs in Drosophila melanogaster [31]. Previous studies showed that, when administered to flies via their food, pharmacochaperones can restore delivery of folding-deficient mutants of DAT to the presynaptic specializations of dopaminergic neurons *in vivo* [32,33]. Accordingly, we examined the effect of ECSI#6 and the reference compound noribogaine on transporter trafficking by generating transgenic flies. Serotonergic neurons innervate the dorsal fan-shaped body (FB) neuropil in the central brain of *Drosophila melanogaster*. These projections can be visualized by expressing membrane-anchored GFP (i.e. GFP fused to the C-terminus of murine CD8; [34]) under the control of TRH-T2A-Gal4 (Fig. 6, top panel). Similarly, when placed under the control of TRH-T2A-Gal4, YFP-tagged wild-type human SERT was delivered to the fan-shaped body (Fig. 6, lefthand image in the middle row). In contrast, in flies harboring human SERT-PG^601,602^AA, the fan-shaped body was devoid of any specific fluorescence (Fig. 6, left-hand image in the bottom row). However, if three-day old male flies expressing human SERT-PG^601,602^AA were fed with food pellets containing 100 μM ECSI#6 or 100 μM noribogaine (Fig. 6 right-hand image in the bottom row) for 48 h, fluorescence accumulated to a level, which allowed for delineating the fan-shaped body (Fig. 6, middle and right-hand image in the bottom row, respectively). This show that ECSI#6 and noribogaine exerted a pharmacochaperoning action *in vivo*, which partially restored the delivery of the mutant transporter to the presynaptic territory. As expected, in flies harboring wild-type human SERT, feeding of ECSI#6 and noribogaine did not have any appreciable effect on the level of fluorescence in the fan-shaped body (Fig. 6, middle and right-hand image in the middle row, respectively).

**Fig 6.**
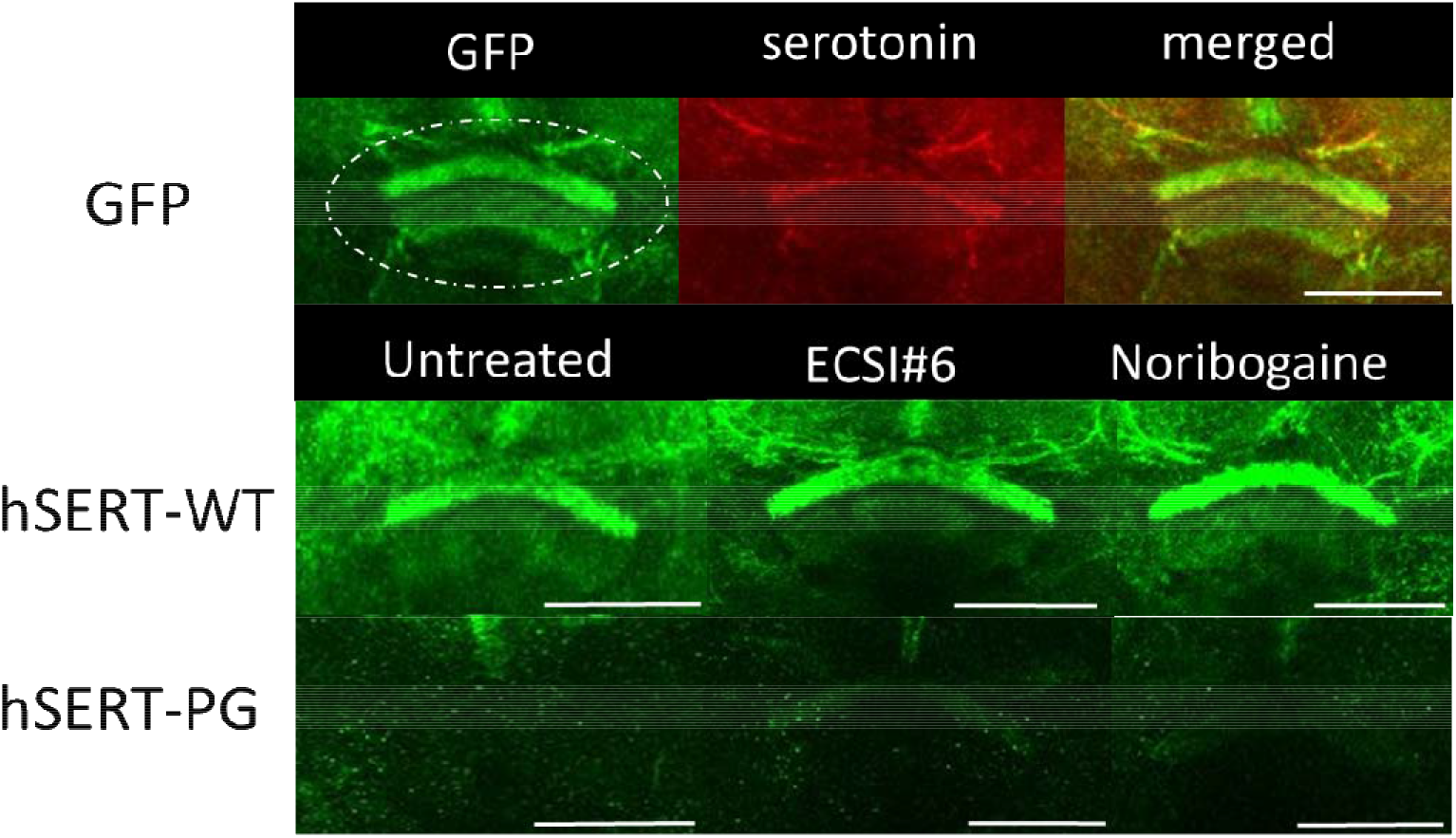
ECSI#6 and noribogaine modulate presynaptic expression of hSERT-PG^601,602^AA in the adult fly brain. The top row shows the expression of membrane-anchored GFP in the presynaptic compartment of the fan-shaped body (denoted by the dotted circle, *UAS-mCD8GFP; TRH-T2A-GAL4;*), labeling with an anti-serotonin antibody (red) and the merged image. The middle and bottom row show the expression of YFP-tagged human wild-type SERT (hSERT-WT *;;UAS-YFP-hSERT-WT/ TRH-T2A-GAL4;)* and SERT-PG^601,602^AA (hSERT-PG *;;UAS-YFP-hSERT-PG^601,602^AA/TRH-T2A-GAL4*;), respectively, in the fan-shaped body. Three-day old male flies expressing hSERT-WT and hSERT-PG were treated with 100 μM ECSI#6 (middle column) and 100 μM noribogaine (right column) for 48h. Images from the brains of these flies were captured by confocal microscopy and compiled with the ImageJ software. Images are representative of at least 10 brains per condition (scale bar□=□50□μm).

### Cocaine, noribogaine and ECSI#6 bind preferentially to different conformations of SERT

We implemented a simple kinetic model that represents the transport cycle of SERT (reaction scheme in black, Fig. 7A) to recapitulate the experimental observations on transport inhibition by cocaine, noribogaine and ECSI#6. In the model, the transport cycle begins with the binding of sodium (Na^+^) and substrate (S) to SERT to the apo state (To). This causes the transporter to isomerize from the substrate-bound outward open (ToNaS) to the substrate-bound inward open (TiNaS) state. Subsequently, Na^+^ and substrate dissociate on the intracellular side and SERT completes the cycle by anti-porting K^+^ (TiK -> ToK). The reactions outlined in grey in Fig. 7A represent the possible states of SERT to which an inhibitor can bind.

**Fig 7.**
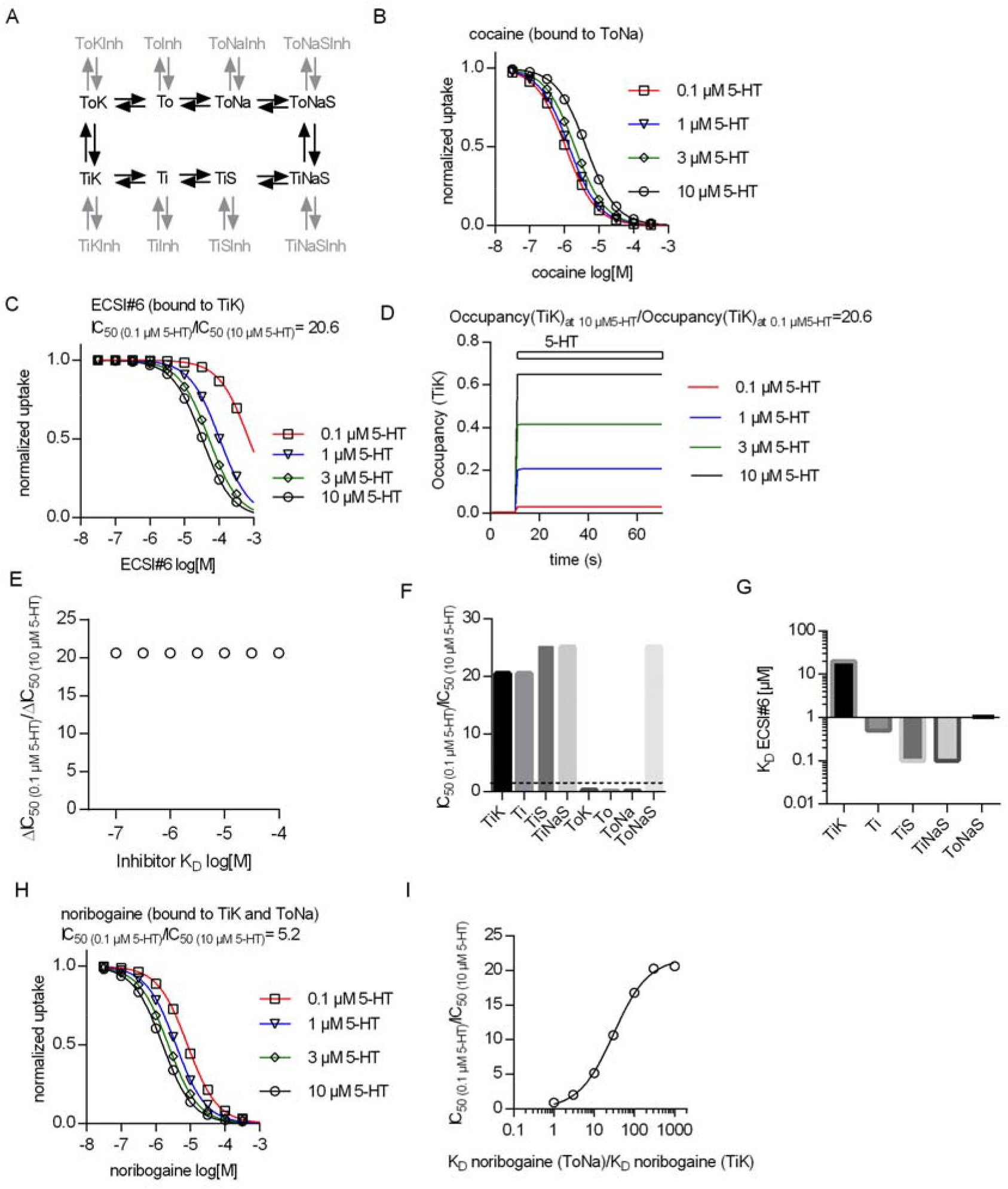
Conformational preference of cocaine, noribogaine and ECSI#6 for binding to SERT. **A,** Reaction scheme (black) of a simple kinetic model of substrate transport by the serotonin transporter (SERT). The chosen microscopic rate constants were as follows: k_on_(K^+^) = 10^6^ M^-1^*s^-1^ k_off_(K^+^) = 5000 s^-1^, k_on_(Na^+^) = 10^6^ M^-1^*s^-1^, k_off_(Na^+^) = 1000 s^-1^, k_on_(5-HT) = 10^7^ M^-1^*s^-1^, k_off_(5-HT) = 500 s^-1^, ToNaS TiNaS = 60 s^-1^, TiNaS ToNaS = 75 s^-1^, TiK ToK = 5 s^-1^, ToK→TiK = 4 s^-1^. The reactions in grey represent possible states where a SERT-specific inhibitor can bind. **B,** Simulation of uptake inhibition by cocaine matches experimental data when the preferred state of cocaine binding is to the ToNa state. **C,** Simulation of uptake inhibition by ECSI#6 matches experimental data when the preferred state of ECSI#6 binding is to the TiK state. The ratio of ECSI#6 IC_50_ in the presence of 0.1 and 10 μM 5-HT (IC_50(10)_/IC_50(10)_) is equal to 20.6. **D,** Simulations of occupancy of TiK states with increasing concentrations of 5-HT over time. The ratio of steady-state 5-HT occupancy of TiK at 10 μM and 0.1 μM is also 20.6. **E,** The observed shift in the ratio IC_50(10)_/IC_50(10)_ is not affected, if the affinity of ECSI#6 for SERT is varied. **F,** Assessment of the IC_50(0.1/10)_ ratio upon assigning binding preference of ECSI#6 to different possible inhibitor bound states. ECSI#6 shows a clear preference for the inward-facing states of SERT. The notable exception is the ToNaS state, which requires binding to an allosteric site in SERT. **G,** The true affinity estimates of ECSI#6 were extracted from the simulations by assuming preferential binding to the indicated distinct states of SERT. **H,** Simulation of uptake inhibition by noribogaine matches the experimental data, if the preferred state of noribogaine binding is to both ToNa and TiK states. **I**, A ratio of IC_50(10)_/ IC_50(10)_ of 5, which was close to that observed in the actual experiments with noribogaine, can be recapitulated in the simulations, if the K_D_ of noribogaine to the ToNa state is assumed to be 10-fold higher than the K_D_ to the TiK state.

Inhibition of substrate uptake by cocaine can be modeled as competitive inhibition, where the inhibitor precludes binding of 5-HT to SERT in the ToNa state (Fig. 7B). In contrast, it is necessary to posit preferential binding of ECSI#6 to the TiK state of SERT for recapitulating the experimental observations by the synthetic data (cf. Fig. 1D and Fig. 7C). We extracted from these simulations the IC_50_-values for uptake inhibition by ECSI#6 in the presence of 0.1 μM and 10 μM 5-HT: the ratio IC_50(0.1)_/IC_50(10)_ was 20.6, which is reasonably close to the shift of 17.2 observed in Fig. 1D. This value matches the ratio of occupancy of the TiK state at 10 and 0.1 μM 5-HT (Fig. 7D). Thus, increasing 5-HT concentrations raise the availability of SERT in the TiK state and this - rather than the affinity of ECSI#6 - accounts for the enhanced ability of ECSI#6 to block 5-HT transport. This interpretation was further substantiated by simulations, where the affinity of ECSI#6 to the TiK state was varied: a variation of the K_D_ over three orders of magnitude did not alter the IC_50(0.1)_/IC_50(10)_ ratio (Fig. 7E). Interestingly the IC_50(0.1)_/IC_50(10)_ ratios remain high, if ECSI#6 binding is assumed to have preference to other inward open transporter states (i.e., Ti, TiS and TiNaS). As expected, if ECSI#6 is assumed to bind preferentially to the outward-open states (i.e., ToK, To, and ToNa), the IC_50(0.1)_/IC_50(10)_ ratios decrease below 1 (Fig. 7F). An exception is the high IC_50(0.1)_/IC_50(10)_ ratio, which is observed, if ECSI#6 binding is assumed to the ToNaS state. This, however, requires binding to an allosteric site, which is unlikely: none of our experimental observations support occupancy of this state by ECSI#6. Thus, the most plausible explanation is binding of ECSI#6 to the TiK state. We estimated the binding affinity of ECSI#6 for TiK from the synthetic data (Fig. 7G): a K_D_ of 12 μM sufficed to reproduce all experimental data. In contrast, the experimental data were only emulated, if the binding affinities to the other preferred states (i.e., TiS, TiS, TiNaS and ToNaS, cf. Fig. 7F) were much higher, that is, in the submicromolar range for TiS, TiS, TiNaS or 1 μM for ToNaS (Fig. 7G). Thus, binding of ECSI#6 to these states provides a possible, but highly unlikely solution.

Finally, we also interrogated the model to account for the shift in the concentration-response curves for uptake inhibition by noribogaine (*cf.* Fig. 1C); the shift was less pronounced than that seen for ECSI#6. These observations can be accounted for in the simulations (Fig. 7H) by assuming preferential binding of noribogaine to both, the ToNa and the TiK states of SERT. If K_D_ for ToNa is assumed to be 10 times larger than the K_D_ for TiK, the kinetic model simulations yield a IC_50(0.1)_/IC_50(10)_ ratio of 5 for noribogaine. This ratio is reasonably close to the ratio of 3, which was observed in the actual experiments. We also systematically varied the ratio of K_D_(ToNa)/K_D_(TiK) to examine the ratio required to reach a ratio IC_50(0.1)_/IC_50(10)_ of 17 to 21, i.e. the range seen with ECSI#6. As evident from Fig. 7I, the K_D_(ToNa) must be 100 to 1000-fold larger than K_D_(TiK). Hence, based on these calculations, it is appropriate to conclude that ECSI#6 has a much higher preference for binding to TiK than noribogaine. In contrast, the binding mode of noribogaine is mixed. This accounts for the observation that the phenotypic consequence of noribogaine binding is a type of uptake inhibition, which is intermediate between that of cocaine and that of ECSI#6.

We verified this interpretation by simulating the saturation kinetics of 5-HT transport by SERT in the presence of the inhibitor. For noribogaine, the simulations show a reduction in the V_max_ of 5-HT transport with increasing noribogaine concentrations (Fig. 8A); the concentration-dependent drop in V_max_ was in reasonable agreement with the experimental observations (cf. Fig. 3B). However, if the K_M_ of 5-HT transport by SERT was plotted as a function of noribogaine concentration, the analysis revealed an initial drop in K_M_, which leveled off (Fig. 8C). This finding is consistent with binding of noribogaine to both, ToNa and TiK state of SERT. The initial drop in K_M_ is too small to allow for its detection within the experimental inter-assay variation of K_M_ estimates (*cf.* Fig. 3E), while the reduction in V_max_ is readily seen (*cf.* Fig. 3H). Hence, based on experimental observations, uptake inhibition by noribogaine is classified as non-competitive. We stress, however, that this neglects the complex binding mode of noribogaine. In contrast, the simulations of substrate transport in the presence of ECSI#6 recapitulated the reduction in the K_M_ and V_max_ of 5-HT transport by SERT (Fig. 8B and D), which was seen in the actual experiments summarized in Fig. 3C, F and I, and thus confirmed the uncompetitive mode SERT of inhibition.

**Fig 8.**
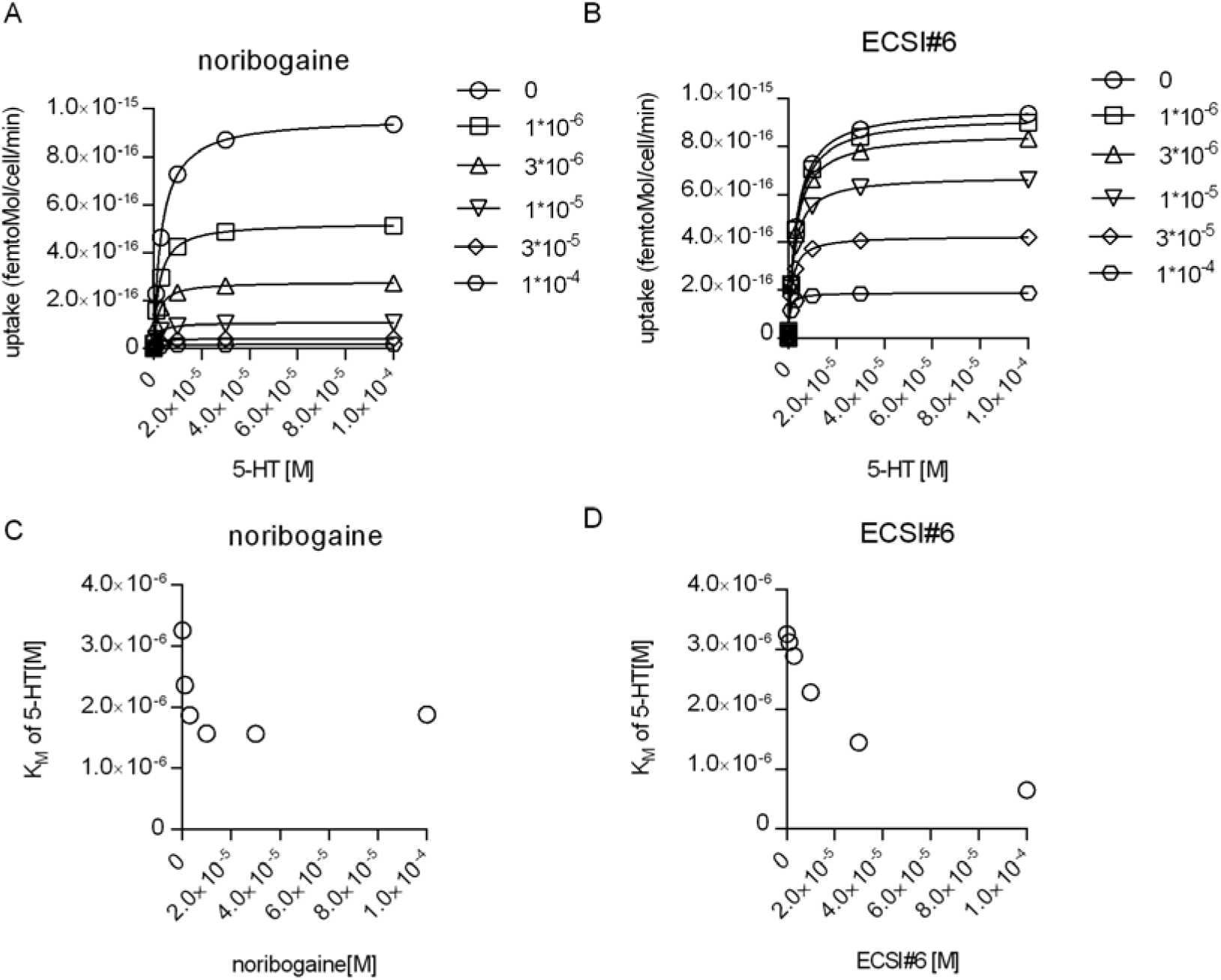
Interrogating the kinetic model to analyze the inhibition of substrate transport by SERT by noribogaine and ECSI#6. **A & B,** Simulations of the saturation kinetics of SERT in the presence of the indicated concentrations of noribogaine **(A)** or ECSI#6 **(B)**. The curves were generated using the reaction scheme depicted in Fig. 7A and the parameters calculated in Fig. 7 for noribogaine and ECSI#6. The K_M_ of 5-HT uptake by SERT was extracted by fitting the data points to a rectangular hyperbola and plotted as a function of the concentration of noribogaine **(C)** and ECSI#6 (**D**).

## Discussion

SERT is arguably the best understood SLC transporter [35]. Atomic details are available for several conformations of SERT [36–38]. The ligand binding pocket and its coupling to the vestibular binding site have been probed by atomic force microscopy, which defined the energy landscape of the binding interaction at the single molecule level (39,40). The kinetics of its transport cycle have been extracted from recordings in real-time, which also allowed for deducing the cooperativity of substrate and co-substrate binding [2,8,13,18,23,24]. The rich pharmacology of SERT comprises typical and atypical inhibitors, partial and full releasers and allosteric modulators [8,10,11]. Here, we identify ECSI#6 as a SERT inhibitor with unique properties that are not matched by any of the known ligands: Substrate enhanced the ability of ECSI#6 to inhibit transport resulting in a progressive lowering of the IC_50_ with increasing concentrations of substrate. In a reciprocal manner, ECSI#6 decreased the K_M_ and the V_max_ of SERT for substrate translocation. This is the hallmark of uncompetitive inhibition. To the best of our knowledge, this is the first report of an uncompetitive inhibitor of SERT and, in fact, of any transporter. Uncompetitive inhibition is rare [41]. In enzymes, uncompetitive inhibition requires binding of the inhibitor to the enzyme-substrate complex [41]. Accordingly, the kinetic model also offered allosteric binding of ECSI#6 to the fully loaded outward-facing transporter - i.e. the ternary complex ToNaS of SERT, substrate and sodium - as one possible solution. However, several arguments rule out this solution: (i) The simulations showed that ECSI#6 must bind with K_D_ of 1 μM to ToNaS to recapitulate the experimental data. However, in the presence of subsaturating 5-HT (10 μM), the affinity estimate of ECSI#6 was >30-fold lower (*cf*. Fig. 2C). (ii) It is also evident from our observations that ECSI#6 and serotonin were not bound simultaneously to SERT: raising the serotonin concentration shifted the curves for displacement of [^3^H]citalopram by ECSI#6 to the right. Thus, ECSI#6 binds to the same site as serotonin, albeit in a different mode: binding of ECSI#6 was favored in the presence of potassium. The resulting affinity estimate (5 μM, *cf.* Fig. 2F) was in the range of the KD-values for the inward-facing potassium bound conformation of SERT TiK required by the kinetic model to recapitulate the experimental data. (iii) The electrophysiological recordings showed that the block of the steady-state, transport-associated current was imposed by ECSI#6 upon binding to an inward-facing state of SERT. (iv) This conclusion was also confirmed by the other solutions provided by the kinetic model: in these solutions, uncompetitive inhibition by ECSI#6 was accounted for by binding of ECSI#6 to the inward-facing states of SERT. We note that an allosteric action of ECSI#6 - i.e. binding to the inward-facing ternary complex TiNaS - was also permitted by the model. However, we consider this unlikely a realistic solution, because - as already pointed out - binding of ECSI#6 and serotonin was mutually exclusive. Hence, we do not consider the analogy of an enzyme-substrate complex, which recruits the inhibitor [41], appropriate to understand the mechanisms underlying uncompetitive inhibition of SERT by ECSI#6. In ligand-gated ion channels, uncompetitive inhibition is observed with open channel blockers. This is exemplified by the NMDA-receptor, where dizocilpine maleate/MK-801, memantine, ketamine and related compounds require the presence of agonists to reach their binding site and to establish blockage of the channel [42]. This use-dependent inhibition is more useful as a starting point than the enzyme-substrate complex to understand the mechanisms underlying uncompetitive inhibition of SERT by ECSI#6: in the presence of physiological ion gradients, SERT accumulates in the outward-facing state [23]. Serotonin is required to drive SERT into the transport cycle, where SERT accumulates in the inward-facing state, because the return step (TiK -> ToK, cf. Fig. 7A) is rate-limiting [23]. This causes the inhibitory action of ECSI#6 to be use-dependent, i.e., enhanced by substrate. Thus, preferential binding of ECSI#6 to the potassium-bound, inward-facing state of SERT (TiK) is sufficient to account for all experimental data.

Our detailed analysis also shed light on the enigmatic properties of ibogaine. Ibogaine and its more potent analog noribogaine have been classified as non-competitive inhibitors. This mode of inhibition was attributed to their trapping the transporter in the inward-facing conformation [20,21]. In fact, the structure of the complex of ibogaine and SERT reveals an inward-facing conformation [38]. Surprisingly, however, ibogaine binds to SERT from the extracellular side; it cannot gain access to SERT via the inner vestibule [25]. Access from the extracellular side also explains, why ibogaine and carbamazepine, which binds to the vestibular (S2) site, can be bound concomitantly [17]. Finally, sole binding of ibogaine or noribogaine to the inward-facing conformations ought to cause uncompetitive inhibition in a manner similar to ECSI#6. The current observations solve this conundrum: the kinetic model showed that an interaction of noribogaine with both, the sodium-bound, outward-facing state (ToNa) and the potassium-bound, inward-facing state (TiK) of SERT was required to recapitulate the experimental observations. The kinetic model also revealed the predicted uncompetitive component, which was, however, too small to be detectable in the experimental noise. It is worth pointing out that the overall K_M_ of substrate transport is a compound parameter, which is affected by both, the rates of individual partial reactions and the occupancy of the individual states in the transport cycle [43,44]. Based on the insights provided by the combination of experimental observations and kinetic modeling, we conclude that, in contrast to ECSI#6, noribogaine has a mixed binding mode.

The endpoint of the folding trajectory of a protein is a minimum energy conformation. The outward and the inward-facing conformation are equivalent stable folds [45]. The ionic conditions in the endoplasmic reticulum (ER) differ from those at the cell surface. The absence of a sodium gradient favors the inward-facing conformation of SERT and other SLC6 transporters. Thus, it is reasonable to posit that their folding trajectory visits the inward-facing conformation [26,27]. This conjecture is supported by the observation that mutations, which trap SERT in the inward-facing state, act as second site suppressors: they restore export from the endoplasmic reticulum and cell surface delivery of folding-deficient mutants [46]. In addition, many compounds, which act as pharmacochaperones on SERT or DAT, are atypical inhibitors with appreciable affinity for the inward-facing state [11,12,28,30,32,33,47,48]. Thus, based on its preferential binding to the inward-facing state of SERT, ECSI#6 was predicted to have pharmacochaperoning activity. This prediction was verified with SERT-PG^601,602^AA: pretreatment with ECSI#6 corrected the folding defect of this mutant, because it restored export from the endoplasmic reticulum and substrate transport by SERT-PG^60,02^ AA. In addition, administration of ECSI#6 to flies resulted in delivery of the mutants to the presynaptic specialization of the axonal projection to the fan-shaped body. We note, though, that noribogaine was more efficacious as a pharmacochaperone: Emax of pharmacochaperoning, i.e., the extent to which substrate transport was restored at saturating concentrations - was consistently higher with noribogaine than with ECSI#6. Variable efficacy of pharmacochaperones was noted previously [30,48]. Efficacy also depends on the nature of the mutation: individual mutants are stalled at different positions within the energy landscape of the folding trajectory [27–29,46]. Thus, individual mutants must differ in their susceptibility to pharmacochaperoning. In addition, in a given folding-deficient mutant, closely related compounds differ in E_max_ of pharmacochaperoning [13,30,48]. Thus, pharmacochaperoning efficacy is also an intrinsic property of a compound linked to its chemical structure. Presumably, for a given mutant, variation in pharmacochaperoning efficacy reflects the different ability of individual compounds to stabilize one or several folding intermediates. There are four mechanisms, which account for pharmacochaperoning [49]: (i) binding to and stabilization of the folded state shifts the folding equilibrium and prevents the backward reaction; (ii) binding to folding intermediates smoothens the energy landscape and precludes stalling of the trajectory in unproductive traps of local minima; (iii) prevention of aggregate formation and (iv) dissolution of aggregates can maintain or restore the folded state. The latter two mechanisms are immaterial to SERT: misfolded SERT does not form aggregates in the endoplasmic reticulum [50]. The EC_50_ of ECSI#6 for pharmacochaperoning was lower than the concentration required for half-maximum inhibition of substrate uptake at saturating substrate. This is circumstantial evidence for an action of ECSI#6 on folding intermediates. Regardless of the underlying mechanism, because ECSI#6 was more potent a pharmacochaperone than a transport inhibitor, it is an attractive starting point to search for more efficacious compounds with pharmacochaperoning activity. In addition, *in vivo*, the action of ECSI#6 on serotonergic transmission is likely to differ from typical competitive inhibitors: because of the uncompetitive mode of its inhibition, ECSI#6 is predicted to cause a use-dependent block of SERT. Serotonergic pathways, which when activated, are more susceptible to modulation by ECSI#6. The reverse is true for competitive - i.e., surmountable - inhibitors.

## Supplementary Materials

not applicable

## Acknowledgments

We thank Sándor Antus (University of Debrecen, Hungary) for the generous gift of ECSI#6.

## Author Contributions

Conceptualization, S.B., C.P., M.F. and W.S.; methodology, S.B., A.E.-K., A.K., D.B., S.S. and W.S.; software, W.S.; validation, S.B., T.H., M.F. and W.S.; formal analysis, S.B., A.H., A.E.-K., A.K., M.F. and W.S.; investigation, S.B., A.E.-K., A.K., D.B., J.B.R.; resources, T.H., S.S., M.F. and W.S.; data curation, M.F. and W.S.; writing—original draft preparation, S.B. and M.F.; writing—review and editing, M.F. and W.S.; visualization, S.B., A.E.-K., A.K. and W.S.; supervision, M.F. and W.S.; project administration, W.S.; funding acquisition, S.S., M.F. and W.S. All authors have read and agreed to the published version of the manuscript

## Institutional Review Board Statement

not applicable

## Informed Consent Statement

not applicable

## Data Availability Statement

All data are contained within the manuscript. Original data are available upon reasonable request.

